# Melatonin alleviates acid-induced stress and improves peanut (*Arachis hypogaea* L.) growth in a dose-dependent manner

**DOI:** 10.64898/2026.01.29.700749

**Authors:** Muhammad Hafeez Ullah Khan, Ruoyi Fu, Ali Muhammad, Shuaichao Zheng, Daijing Zhang, Zhiyong Zhang, Quanyong Liu

**Affiliations:** Xinxiang Key Laboratory of Crop Root Biology and Green Efficient Production/Henan Key Laboratory of Quality and Stress-Resilient Bio-breeding and Safe Production of Cotton and Oil Crops, School of Life Sciences, Henan Institute of Science and Technology, Xinxiang, Henan, 453000, China; College of Life Sciences, Henan Normal University, Xinxiang 453007, Henan, China

**Keywords:** Melatonin supplementation, Peanut seedling resilience, Acidic stress, ROS-pH coordination, Low-pH crop physiology

## Abstract

Acidic stress severely restricts crop growth by disrupting nutrient uptake, redox homeostasis, and membrane stability, yet mitigation strategies remain limited. Here, we investigated the role of melatonin (MT) in regulating growth, photosynthesis, oxidative stress, antioxidant defense and proton transport in peanut seedlings under controlled hydroponics acidic (pH 4.0) and near-optimal (pH 6.5) conditions, and validated these findings in naturally acidic field soil (pH 4.3-4.5). Acid stress markedly reduced biomass accumulation, chlorophyll content, and redox balance, while enhancing ROS (H_2_O_2_) and lipid peroxidation (MDA). Exogenous MT application, particularly at 50-100 µM, significantly improved shoot and root biomass, restored chlorophyll pigments and reduced H_2_O_2_ and MDA accumulation, with more pronounced effects under pH 4.0 than pH 6.5. MT strongly activated antioxidant enzymes (SOD, CAT, APX), while POD activity declined, reflecting melatonin’s dual role as both a direct ROS scavenger and a regulator of enzymatic redox networks. Notably, MT induced strong, dose-dependent upregulation of HL-ATPase genes (AH1 and AH2) in both leaves and roots under acidic conditions, suggesting enhanced proton extrusion, intracellular pH homeostasis, and stress adaptation. The soil validation experiment confirmed the agronomic relevance of these findings, where MT dose-dependent concentrations improved germination, vegetative growth, chlorophyll fluorescence (Fv/Fm), and yield-related traits under natural acidic conditions. Although MT also conferred benefits at pH 6.5, responses were generally moderate compared with acid stress. Collectively, these results demonstrate that MT enhances peanut tolerance to acid stress across both controlled and natural field-relevant environments, highlighting its potential application for sustainable crop production on low-pH soils.

## 1. Introduction

Soil acidification has become an escalating global concern, now affecting nearly 30–40% of the world’s arable lands and posing a serious threat to agricultural productivity and food security (Huang *et al*., 2023). Soils with a pH below 5.5 are considered acidic, and such conditions limit the availability of essential macronutrients like phosphorus (P), calcium (Ca²□), and magnesium (Mg²□). At the same time, they increase the solubility of toxic metal ions, especially aluminium (Al³□) and manganese (Mn²□), which can be highly detrimental to plant growth (Delfim *et al*., 2025). This imbalance in nutrient and ion dynamics often results in inhibited root elongation, reduced nutrient uptake, and disrupted photosynthetic processes. Furthermore, an excess of hydrogen ions (H□) in the rhizosphere disturbs pH stability and alters microbial community structure, compounding physiological stress in plants (Naz *et al*., 2022; Thapa *et al*., 2025).

Peanut (*Arachis hypogaea* L.), a major legume widely cultivated in tropical and subtropical regions, is particularly vulnerable to acid soil stress (Li *et al*., 2024b; Jamir *et al*., 2025). During early seedling stages, low pH and other abiotic stress conditions restrict root development, damage cell membranes, and interfere with ion transport mechanisms, ultimately impairing water and nutrient absorption. These physiological disruptions often lead to reduced chlorophyll levels, decreased photosynthetic efficiency, and poor biomass accumulation (Liu *et al*., 2018; Borhannuddin Bhuyan *et al*., 2019; Li *et al*., 2021). On a cellular scale, acid stress intensifies the generation of reactive oxygen species (ROS), including superoxide anions (O□□), hydrogen peroxide (H□O□), and hydroxyl radicals (•OH), which can damage lipids, proteins, and nucleic acids (Ahmad *et al*., 2023b; Hasan *et al*., 2025). Although plants possess both enzymatic antioxidants (such as SOD, POD, APX, and CAT) and non-enzymatic compounds (like ascorbate, glutathione, and flavonoids). These defense mechanisms are often inadequate to neutralize the rapid oxidative surge induced by acidic environments, particularly in sensitive crops like peanut.

In recent years, melatonin (N-acetyl-5-methoxytryptamine) has been identified as an effective plant growth regulator and stress alleviator (Zheng *et al*., 2025; Muhammad *et al*., 2026). Melatonin was first found in animals, and is now widely recognized in plants to modulate plant growth, development and defence responses (Ahmad *et al*., 2023a). Exogenous melatonin has been reported to alleviate various abiotic stresses such as drought, salinity, acid, extreme temperature, nutrient deficiency and heavy metal toxicity by enhancing photosynthetic efficiency, maintaining redox balance and elevating antioxidant capacity (Basharat *et al*., 2025; Dou *et al*., 2025; Kong *et al*., 2025). Melatonin is also involved in alleviating aluminium toxicity under suppressed soil pH, maintaining the integrity of cellular membrane, stabilizing chloroplast structure and activating the antioxidant system (Wang *et al*., 2025a). In addition to its direct ROS-scavenging ability, melatonin indirectly regulates redox homeostasis by modulating the expression of redox-responsive genes (Hasan et al., 2025; Jiang *et al*., 2025a).

Proton pumps, particularly plasma membrane and vacuolar H□-ATPases, are fundamental molecular systems that enable plants to sustain cytoplasmic pH homeostasis under an acidic environment (Cosse & Seidel, 2021; Zeng *et al*., 2024; MacDonald *et al*., 2025). These ATPases actively transport protons from the cytoplasm into the apoplast or vacuole, thereby establishing electrochemical gradients that drive essential physiological processes such as nutrient uptake, ion transport, stomatal regulation and stress adaptation (MacDonald et al., 2025; Sun *et al*., 2025). In peanut, various ATPase genes have been characterized, among which AH1 and AH2 are key regulators of intracellular pH balance and ion homeostasis under acid stress. The upregulation of H□-ATPase genes has been reported in several crops as an adaptive mechanism to low pH, facilitating proton extrusion and protecting cellular metabolism (Li *et al*., 2022; Li *et al*., 2024a). However, the regulatory dynamics of these genes under melatonin treatment remain largely unexplored in the peanut seedling.

Melatonin has been shown in other crop species to influence proton pump activity and ion transport. For instance, melatonin-induced activation of H□-ATPases can enhance nutrient absorption, stabilize membrane potential, and promote root development under stress conditions (Huang *et al*., 2022; Tahjib-Ul-Arif *et al*., 2025). Therefore, integrating physiological, biochemical and transcriptional analysis, particularly through expression profiling of AH1 and AH2, can provide valuable mechanistic insight into how melatonin supports pH regulation and overall homeostasis in peanut seedlings under acidic conditions.

The present study aims to elucidate the integrated physiological, antioxidative, and gene-expression analysis underlying melatonin-mediated acid tolerance in peanut seedlings, with a particular focus on AH1 and AH2 gene regulation. Using controlled hydroponic experiments complemented by validation in naturally acidic field soil, we demonstrate that melatonin enhances peanut tolerance to acid stress by improving growth and photosynthetic performance, strengthening antioxidant defense, and upregulating key H□-ATPase genes to stabilize cellular pH and ion homeostasis. These combined laboratory and soil-based findings provide mechanistic and agronomic evidence that melatonin confers effective acid-stress resilience. Unravelling these findings will contribute to the development of sustainable management practices and genetic improvement strategies for peanut cultivation in acid-affected soils.

## 2. Materials and methods

### 2.1 Hydroponic Acid Stress Experiment

#### 2.1.1 Plant material and experimental design

The experiment was conducted in the greenhouse (25□±□2°C, 60-70% RH, 16/8 h light/dark, 200□μmol□m□²□s□¹) at the Henan Institute of Science and Technology, Xinxiang City, Henan Province, China, under controlled environmental conditions. A paper roll culture system was used in a hydroponic setup to evaluate the effects of melatonin on peanut growth under acidic stress conditions. The experiment followed a randomized complete design (RCD) with eight biological replications to ensure statistical reliability.

Uniform, mature, and healthy seeds of peanut (*Arachis hypogaea* L. cv. Xinbaihua 25) were selected as experimental material. Seeds were surface-sterilized with 1% sodium hypochlorite solution for 5 min and rinsed several times with distilled water to remove any residual disinfectant. To investigate the effects of melatonin on plant growth, photosynthetic pigments, antioxidant responses, and gene expression, the seeds were soaked for 10 hours in melatonin solutions of 0.5 μM, 5 μM, 50 μM, and 100 μM. Seeds soaked in distilled water under identical conditions served as the control (CK).

For each replicate, seven seeds were placed vertically on moistened germination paper to ensure uniform orientation and root emergence. The papers were then carefully rolled and secured with rubber bands to maintain contact and prevent unrolling during incubation. Following treatment, the paper rolls containing the seeds were placed vertically in plastic containers filled with half-strength Hoagland nutrient solution prepared according to the modified composition described by Cai et al. (2021) (Cai *et al*., 2021). Two nutrient pH regimes were maintained: a normal condition with pH 6.5 and an acidic condition with pH 4.0. The pH of each solution was carefully adjusted using 1 M HCl or NaOH and monitored daily to ensure consistent stability throughout the experiment. The lower ends of the paper rolls were immersed approximately 2-3 cm into the solution to facilitate the upward capillary movement of nutrients and water while maintaining adequate aeration around the root zone. To prevent nutrient depletion and pH fluctuations, the nutrient solutions were replaced every two days. During the early germination stage, the paper rolls were kept under dark conditions until radicle emergence to ensure uniform germination. After emergence, seedlings were exposed to normal light conditions for continued growth. After 15 days of cultivation, seedlings were harvested for measurements of growth parameters, photosynthetic pigments, antioxidant enzyme activities, and gene expression analyses.

#### 2.1.2 Plant Germination and Growth Parameters

Seed germination was monitored daily until radicle emergence, which was defined as a visible protrusion of at least 2 mm from the seed coat, and the germination percentage (GP) was calculated at day 7. After 15 days of cultivation, seedlings were carefully removed from the paper rolls and gently rinsed with distilled water to remove residual nutrient solution. Shoot length was measured from the base to the tip of the longest leaf, and root length from the base to the tip of the longest root, using a ruler to the nearest millimeter. Fresh weights of shoots and roots were determined separately after blotting with filter paper to remove surface moisture, and samples were then oven-dried at 70 °C for 72 h or until a constant weight was achieved to determine dry weights. All measurements were performed for each seedling in every replicate, and the mean values of the eight replicates were used for statistical analysis. Data are presented as mean ± standard error (SE) and used for subsequent physiological, biochemical, and molecular analyses.

#### 2.1.3 Photosynthetic Pigment Measurement

Photosynthetic pigments, including chlorophyll a, chlorophyll b, and carotenoids, were extracted from 0.2 g of fresh peanut leaves using 80% (v/v) acetone. The leaf samples were homogenized in chilled acetone, kept on ice for 10-15 min in the dark, and centrifuged at 10,000 × g for 10 min at 4 °C. The clear supernatant was collected, and absorbance was recorded at 663, 645, and 470 nm using a UV–visible spectrophotometer with 80% acetone as blank. Pigment contents were calculated using standard equations and expressed as mg g□¹ fresh weight. All extractions were performed under low-light conditions to minimize pigment degradation, and six independent replicates were analyzed per treatment.

#### 2.1.4 Antioxidant Enzyme Assays

To assess the antioxidant defense response in peanut seedlings, the activities of catalase (CAT), superoxide dismutase (SOD), peroxidase (POD), and ascorbate peroxidase (APX), content, were determined from fresh leaf tissues. Approximately 0.5 g of fresh leaves was homogenized in 5 m□ of ice-cold 50 mM phosphate buffer (pH 7.0) containing 1 mM EDTA and 1% (w/v) polyvinylpyrrolidone (PVP). The homogenate was centrifuged at 12,000 × g for 15 min at 4 °C, and the resulting supernatant was used for all enzyme assays.

Superoxide dismutase (SOD; EC 1.15.1.1) activity was assayed by the inhibition of nitroblue tetrazolium (NBT) reduction under light, with one unit of SOD activity defined as the amount of enzyme required to cause 50% inhibition of NBT photoreduction. Catalase (CAT; EC 1.11.1.6) activity was determined by monitoring the decrease in absorbance at 240 nm due to H□O□ decomposition. Peroxidase (POD; EC 1.11.1.7) activity was measured using guaiacol as a substrate, recording the increase in absorbance at 470 nm as guaiacol was oxidized. Ascorbate peroxidase (APX; EC 1.11.1.11) activity was estimated by monitoring the decrease in absorbance at 290 nm as ascorbate was oxidized in the presence of H_2_O_2_.

#### 2.1.5 Quantification of Malondialdehyde (MDA) and Hydrogen Peroxide (H_2_O_2_)

The levels of malondialdehyde (MDA) and hydrogen peroxide (H_2_O_2_) were determined to evaluate lipid peroxidation and oxidative stress in peanut leaves and roots. Lipid peroxidation was determined by quantifying malondialdehyde (MDA) using the thiobarbituric acid (TBA) reaction. For MDA estimation, 1 mL of enzyme extract was mixed with 2 mL of 0.6% (w/v) TBA in 10% (w/v) trichloroacetic acid (TCA) and heated at 95 °C for 15 min, then rapidly cooled and centrifuged at 10,000 × g for 10 min. The absorbance of the supernatant was measured at 532 nm and corrected for nonspecific turbidity by subtracting absorbance at 600 nm. MDA content was calculated using the extinction coefficient (ε = 155 mM□¹ cm□¹) and expressed as nmol g□¹ fresh weight (FW).

Hydrogen peroxide (H_2_O_2_) content was determined spectrophotometrically using a colorimetric KI reaction with quantification against a standard curve. Fresh leaf tissue (0.5 g) was ground in 5 mL of cold 0.1% (w/v) TCA and centrifuged at 12,000 × g for 15 min at 4 °C. A reaction mixture was prepared by mixing 0.5 mL of the extract with 0.5 mL of 10 mM potassium phosphate buffer (pH 7.0) and 1.0 mL of 1 M potassium iodide (KI) (all kept on ice and protected from light). The reaction was allowed to proceed for 1 min at room temperature in the dark, and the absorbance was read at 390 nm. A standard calibration curve was prepared in parallel using known concentrations of H_2_O_2_ (for example 0, 10, 25, 50, 100, 200 µM) prepared in 0.1% TCA; sample H_2_O_2_ content was interpolated from this curve and expressed as µmol H□O□ g□¹ FW (adjusting for extract and assay volumes and any dilutions). All reactions and standards were run in technical triplicate and blank reactions (reagents without plant extract) were included to correct baseline absorbance.

#### 2.1.6 Gene Expression Analysis

Total RNA was extracted from peanut roots and leaves using the RNAprep Pure Plant Plus Kit (Tiangen Biotech, Beijing, China) according to the manufacturer’s instructions. The quality and quantity of RNA were determined using a NanoDrop 2000 spectrophotometer (Thermo Fisher Scientific, USA) by measuring the absorbance ratios at 260/280 nm and 260/230 nm. RNA integrity was confirmed by 1.0% agarose gel electrophoresis, which showed sharp, intact 28S and 18S rRNA bands without visible degradation. Only high-quality RNA (A260/A280 ratio between 1.9 and 2.1) was used for cDNA synthesis.

First-strand cDNA was synthesized from 50 µg of total RNA using the PrimeScript™ RT reagent kit with gDNA Eraser (Takara, Japan) to eliminate potential genomic DNA contamination. The synthesized cDNA was diluted 10-fold with nuclease-free water and stored at −20 °C until further analysis.

Quantitative real-time PCR (qRT-PCR) was conducted using TB Green® Premix Ex Taq™ II (Takara, Japan) on a QuantStudio™ 5 Real-Time PCR System (Applied Biosystems, USA). Each 20 µL reaction contained 10 µL of 2× SYBR Green master mix, 0.4 µL each of forward and reverse primers (10 µM), 2 µL of diluted cDNA, and 7.2 µL of nuclease-free water. The PCR cycling conditions were as follows: initial denaturation at 95 °C for 30 s, followed by 40 amplification cycles of 95 °C for 5 s and 60 °C for 30 s. A melting curve analysis (60-95 °C) was performed at the end of each run to confirm amplification specificity.

Each sample was analyzed in triplicate, and the mean values of three technical replicates from three independent biological replicates were used for statistical analysis. The relative transcript abundance of target genes was calculated using the 2□ΔΔCt method (Livak and Schmittgen, 2001). The *AhGAPDH* gene was used as an internal reference due to its stable expression across different tissues and treatments. Primer sequences used in the study are listed in Table 1.

**Table 1.**
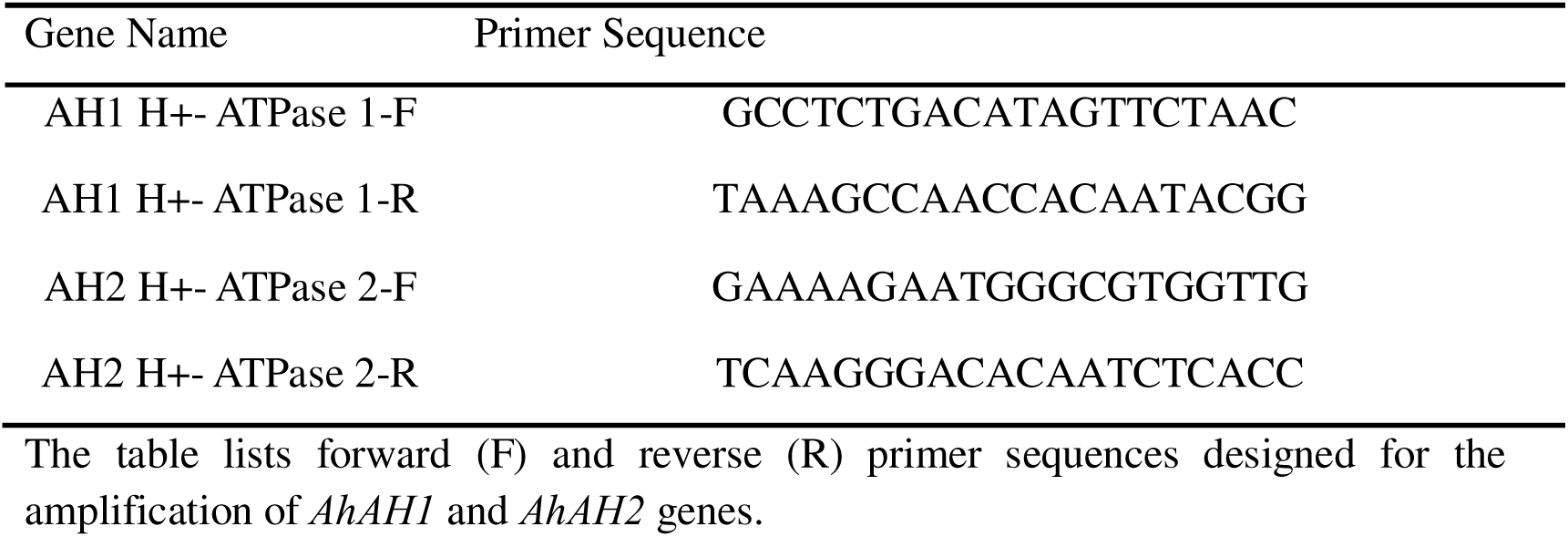
Primer sequences used for qRT-PCR analysis of plasma membrane H□-ATPase genes in peanut.

### 2.2 Field Soil Pot Experiment

#### 2.2.1 Field Soil Collection

Soil samples were collected from agricultural fields located in the southern region of Henan Province, China, where natural soil acidity is commonly observed. Composite soil samples were obtained from the top 0-20 cm layer using a clean stainless-steel auger and transported to the laboratory in sterile polyethylene bags. The soil pH was determined using a soil-to-distilled water ratio of 1:3 and measured with a calibrated digital pH meter, which ranged between 4.3 and 4.5, confirming the naturally acidic condition of the field soil. The collected soil was air-dried, sieved through a 2-mm mesh, and homogenized before pot experimentation.

#### 2.2.2 Field Soil Pot Experiment

Detailed biochemical and molecular analyses were conducted only in the hydroponic system to elucidate mechanistic responses, whereas the soil experiment was designed to validate growth and yield performance under natural acidic conditions. To validate the hydroponic findings under natural soil conditions, a pot experiment was conducted using the collected acidic soil. Peanut seeds (*Arachis hypogaea* L. cv. Xinbaihua 25) were collected and dressed in 0.5 μM, 5 μM, 50 μM, and 100 μM melatonin solution (0.28 g PVA and 0.32 g cellulose were added as an adhesive solution) for 3-5 minutes, which were then air-dried in a cool, shaded place before sowing. Seed dressed only with an adhesive solution, without MT solution, under identical conditions served as the control (CK). Air-dried peanut seeds were sown in plastic pots containing 3 kg of acidic field soil and maintained under natural outdoor conditions from June 2025 to October 2025. The soil field trial was conducted at the experimental field of the Henan Institute of Science and Technology in Xinxiang City, Henan Province, China (114°12′60″ E, 35°21′43″ N), with an average temperature of 28 ± 2□^°^C and a natural photoperiod (≈14 h light/10 h dark). Each treatment consisted of 20 biological replicates with one plant per pot.

#### 2.2.3 Measurement of Growth, Physiological and Yield Traits

Plant growth and physiological parameters were recorded following procedures similar to those described for the hydroponic experiment, with additional yield-related measurements specific to soil cultivation. Germination percentage was calculated at 7 days after sowing. After 15 days, during the vegetative stage, root length, shoot length, fresh weight, and dry weight were measured. Chlorophyll fluorescence (Fv/Fm) was assessed using a portable fluorometer. After 55 days at the maturity stage, reproductive traits, including peg number per plant, pod number per plant, and total pod weight per plant, were recorded to evaluate yield performance under natural acidic soil conditions.

### 2.3 Statistical Analysis

All data were analyzed using SPSS v26.0. Results are presented as mean ± SE. Two-way ANOVA was used to compare treatments, and significant differences were determined using Tukey’s test (p < 0.05). For qRT-PCR, the average Ct values of technical replicates were used, and fold changes were calculated using the 2□ΔΔCt method.

## 3. Results

### 3.1 Effect of melatonin on peanut seed germination under controlled hydroponic acid conditions

Seed germination of peanut was significantly affected by pH and melatonin treatment (Fig. 1a and b). At acidic pH (4), control seeds exhibited the lowest germination (∼70%). Application of MT markedly improved germination in a dose-dependent manner: 0.5 µM MT increased germination to ∼88% (+18%), 5 µM MT to ∼90% (+20%), 50 µM MT to ∼93% (+23%), and 100 µM MT to ∼98% (+28%) with related to control, indicating a strong protective effect of MT under acid stress. Under near-neutral conditions (pH 6.5), germination was consistently high across treatments, with control seeds reaching ∼85% and MT-treated seeds ranging from 90% to nearly 100%. The relative improvement of germination by MT was more pronounced under acidic stress, indicating that MT alleviates pH-induced inhibition. These results demonstrate that melatonin significantly mitigates the inhibitory effects of low pH on peanut seed germination, with the largest relative benefit observed at 100 µM MT under acidic conditions, and that moderate MT concentrations are sufficient to maintain near-maximal germination at near-neutral pH.

**Figure 1.**
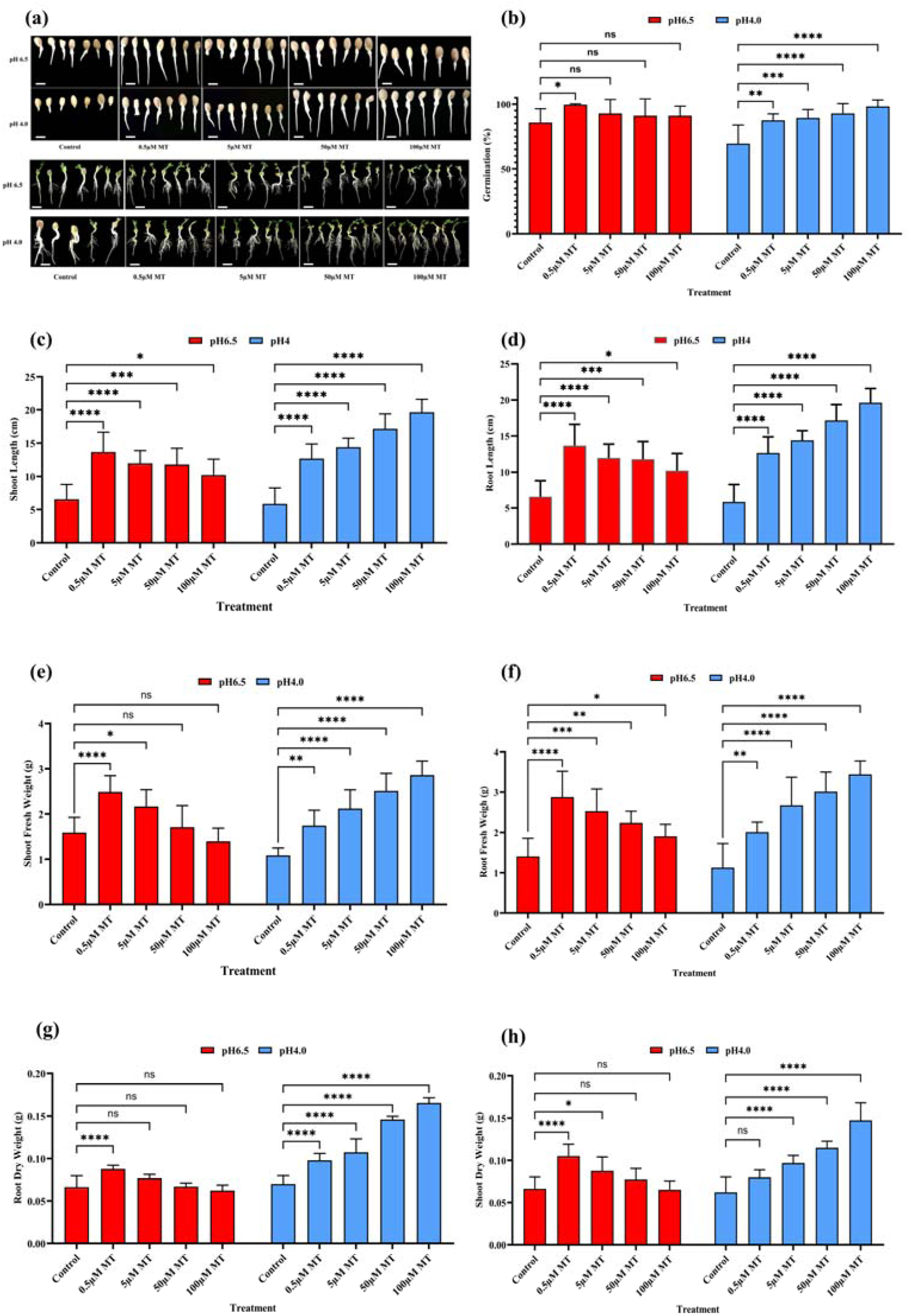
Effect of melatonin (MT) on seed germination and seedling growth under control hydroponic normal (pH 6.5) and acidic (pH 4.0) conditions on peanut. **(a)** seed germination and seedling growth phenotypes, **(b)** seed germination (%), **(c)** shoot length (cm), **(d)** root length (cm), **(e)** shoot fresh weight (g), **(f)** root fresh weight (g), **(g)** root dry weight (g), **(h)** shoot dry weight. Statistical differences among treatments were analyzed using appropriate multiple comparison tests. ns indicates not significant, while *, **, ***, and **** denote significant differences at P < 0.05, P < 0.01, P < 0.001, and P < 0.0001, respectively. Data in the figures are means of eight replications (n = 8) ± standard deviation (SD). Bar = 2 cm.

### 3.2 Melatonin supplementation enhances peanut plant growth and biomass accumulation under controlled hydroponic acid stress

To evaluate the role of melatonin in modulating peanut seedling growth under contrasting pH conditions, root and shoot development as well as biomass accumulation were assessed under acidic (pH 4) and near-neutral (pH 6.5) environments (Figs. 1c–h). Acidic stress (pH 4) markedly suppressed seedling growth, as evidenced by substantial reductions in root and shoot elongation in control plants, indicating strong inhibition of early developmental processes. Melatonin treatment markedly alleviated this inhibition, resulting in a dose-dependent enhancement of root length, shoot length, and biomass. Root length under pH 4 increased progressively from approximately 6 cm in control plants to nearly 20 cm at 100 µM MT, while shoot length increased from about 2.5 cm in the control to over 6 cm at the highest MT concentration, demonstrating a strong stimulatory effect of MT on seedling elongation under acid stress (Fig. 1c and d). Under neutral conditions (pH 6.5), root and shoot lengths were generally higher than those at pH 4 in control plants; however, the growth-promoting effect of MT was less pronounced. Low MT concentration (0.5 µM) enhanced root and shoot length, while higher MT doses showed either moderate improvement or stabilization of growth, suggesting that MT exerts a stronger relative effect under stress conditions than under optimal pH (Fig. 1c and d).

Consistent with growth responses, root fresh weight and shoot fresh weight under pH 4 increased substantially with increasing MT concentration. Root fresh weight rose from 1.09 g in control plants to 3.45 g at 100 µM MT, while shoot fresh weight increased from 1.09 g to 2.86 g, indicating enhanced biomass accumulation (Fig. 1e and f). This trend was further supported by dry weight measurements, where root dry weight increased from 0.07 g in control plants to 0.17 g at 100 µM MT, and shoot dry weight increased from 0.06 g to 0.15 g, confirming that MT-induced growth was associated with true biomass gain rather than water accumulation alone (Fig. 1g and h).

Under pH 6.5, overall growth and biomass parameters were higher than those observed under acidic conditions in the control plants; however, the response pattern to MT differed from that at pH 4. Low MT concentration (0.5 µM) promoted root and shoot growth and fresh weight accumulation, whereas higher MT concentrations (5-100 µM) resulted in a gradual decline in root and shoot length and biomass (Fig. 1e–h). Notably, root dry weight and shoot dry weight at pH 6.5 showed little or no improvement with increasing MT concentration and tended to decrease at higher doses, suggesting a concentration-dependent shift in MT responsiveness under non-stress conditions.

Overall, these results demonstrate that melatonin strongly enhances root and shoot growth and biomass accumulation under acidic stress (pH 4), with maximal effects observed at higher MT concentrations, whereas under near-neutral conditions (pH 6.5), low MT levels are sufficient to promote growth and excessive MT may exert inhibitory effects. This indicates that the growth-promoting role of melatonin is highly dependent on environmental pH and stress intensity.

### 3.3 Melatonin modulates photosynthetic pigments in response to controlled hydroponic acid stress in peanut

Melatonin application significantly influenced the accumulation of photosynthetic pigments in peanut seedlings under both acidic (pH 4) and near-neutral (pH 6.5) conditions (Figs. 2a-d). Under pH 4, control plants exhibited relatively low levels of chlorophyll a, chlorophyll b, total chlorophyll, and carotenoids, indicating impaired pigment biosynthesis under acid stress. However, MT treatments progressively enhanced pigment accumulation, with a clear dose-dependent response.

**Figure 2.**
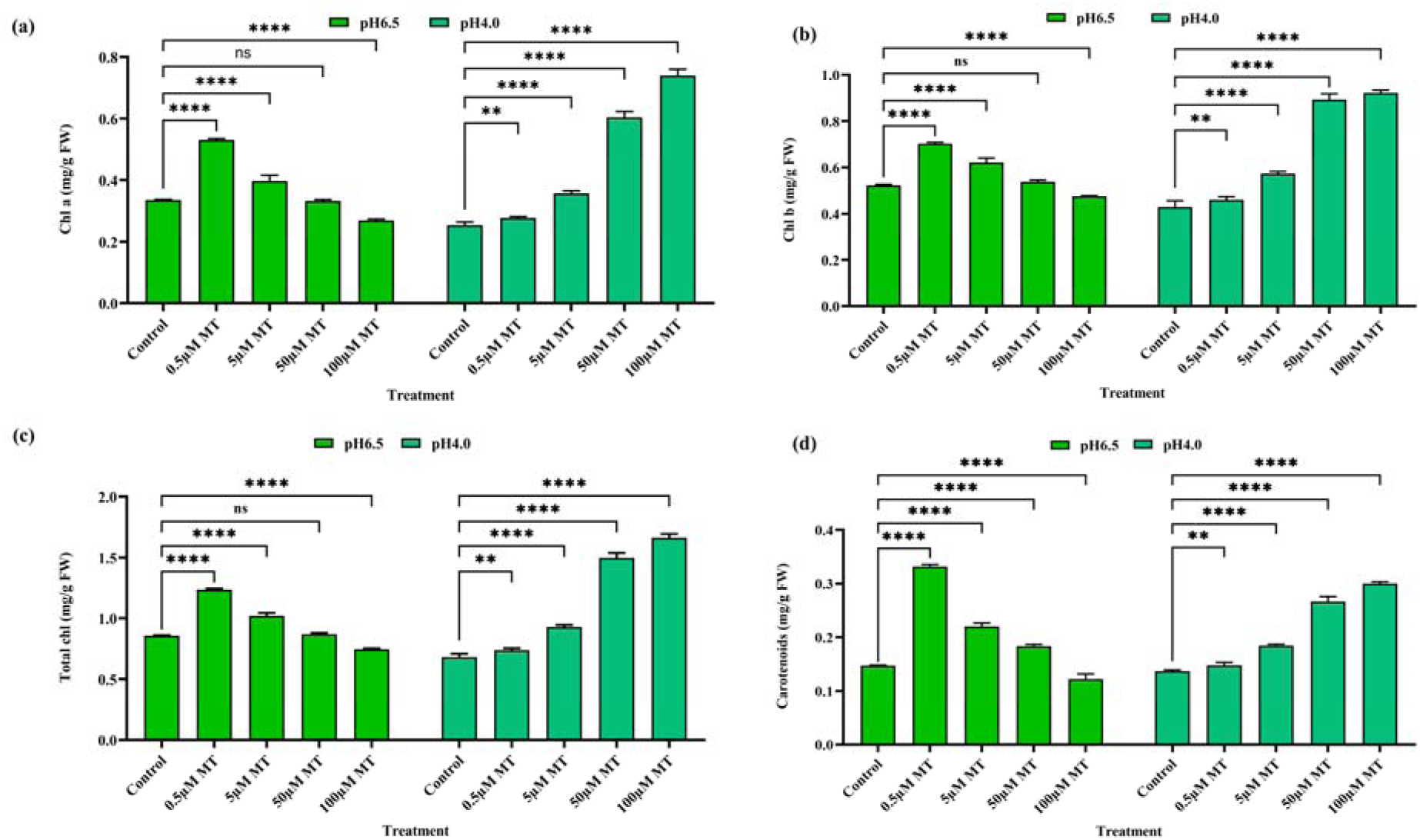
Effect of melatonin (MT) on photosynthetic pigments of peanut under control hydroponic normal (pH 6.5) and acidic (pH 4.0) conditions. **(a)** Chlorophyll *a* (mg/g FW), **(b)** chlorophyll *b* (mg/g FW), **(c)** total chlorophyll content (mg/g FW), **(d)** carotenoids (mg/g FW). Statistical differences among treatments were analyzed using appropriate multiple comparison tests. ns indicates not significant, while ***, and **** denote significant differences at P < 0.001, and P < 0.0001, respectively. Data in the figures are means of eight replications (n = 8) ± standard deviation (SD).

Chlorophyll a content under pH 4 increased gradually with increasing MT concentration, showing moderate enhancement at 5 µM MT and a pronounced increase at 50 and 100 µM MT, where values reached their maximum (Fig. 2a). A similar trend was observed for chlorophyll b, resulting in a substantial rise in total chlorophyll content, particularly at 50 and 100 µM MT. These increases were significantly higher than the control (P < 0.05), demonstrating that MT effectively mitigates acid-induced inhibition of chlorophyll synthesis (Fig. 2b and c). Under pH 6.5, pigment contents were generally higher than those under pH 4 in control plants, reflecting more favorable growth conditions. Notably, low MT concentration (0.5 µM) caused a marked increase in chlorophyll a, chlorophyll b, and total chlorophyll, reaching peak values under this pH. However, higher MT concentrations (50-100 µM) resulted in a gradual decline in pigment levels, suggesting that excessive MT under optimal pH may suppress chlorophyll accumulation (Figs. 2a-c).

Carotenoid content showed a distinct but complementary pattern. Under pH 4, carotenoid levels increased steadily with rising MT concentration, reaching the highest values at 100 µM MT, which were significantly greater than the control (P < 0.05). This indicates enhanced antioxidant capacity under acid stress (Fig. 2d). In contrast, under pH 6.5, carotenoid content peaked sharply at 0.5 µM MT and declined at higher concentrations, paralleling the response observed for chlorophyll pigments (Fig. 2d). Overall, these results demonstrate that melatonin enhances photosynthetic pigment accumulation in peanut seedlings in a pH-dependent and concentration-dependent manner. Low MT concentrations are optimal for pigment enhancement under near-neutral conditions, whereas higher MT doses are more effective in protecting pigment biosynthesis under acidic stress, likely through improved stress tolerance and antioxidant regulation.

### 3.4 Melatonin differentially regulates antioxidant defense system under controlled hydroponic acid stress in peanut

We evaluated the enzymatic antioxidant activity in acid-stressed peanut plants treated with melatonin to understand how exogenous melatonin mitigates acid stress. Our findings revealed that the radical scavenging activities of SOD, POD, CAT, and APX, were markedly affected by both external pH and melatonin treatment in peanut leaves, reflecting dynamic regulation of reactive oxygen species (ROS) homeostasis under stress and non-stress conditions (Figs. 3a-h).

**Figure 3.**
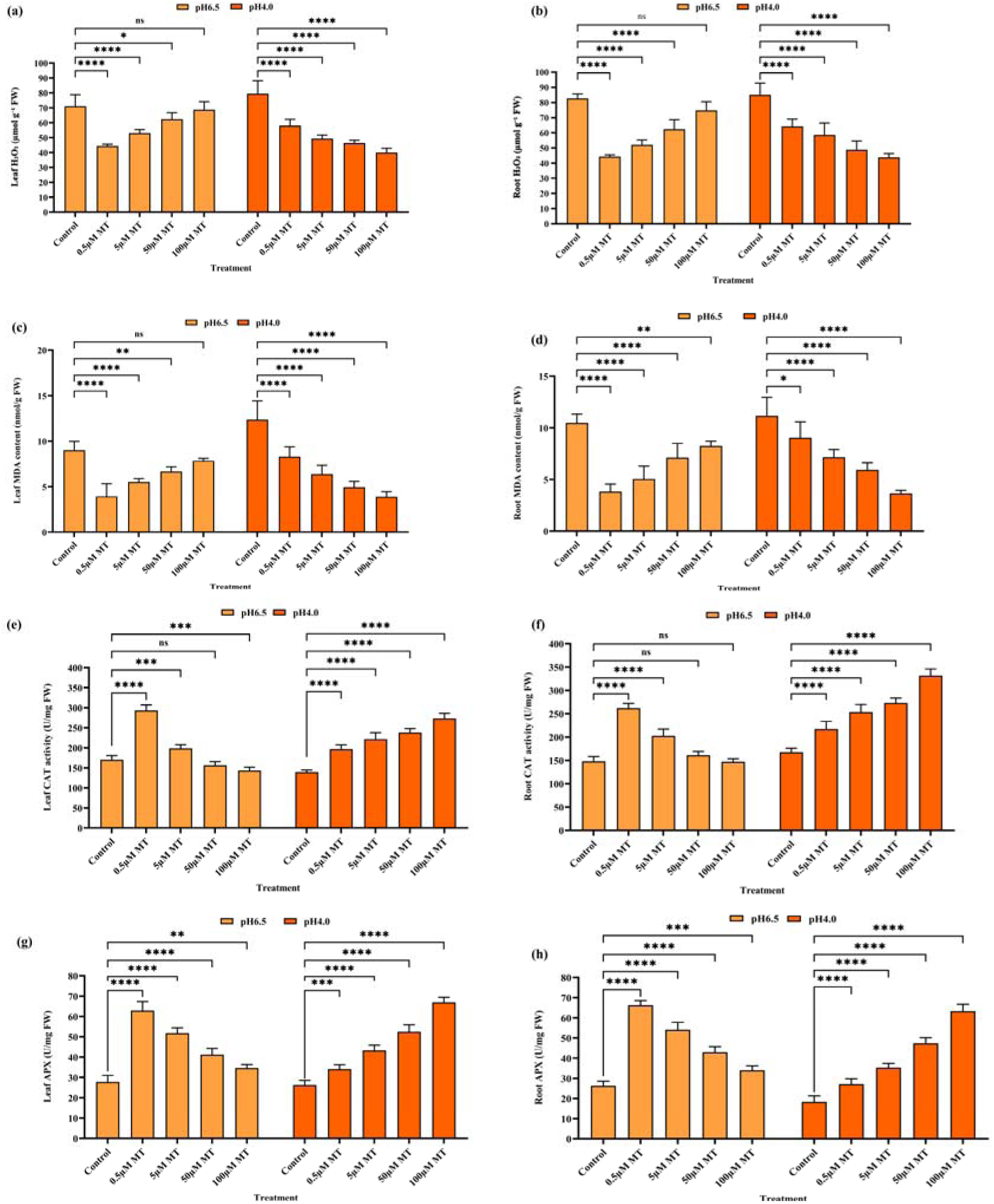
Effect of melatonin (MT) on antioxidant contents of peanut under control hydroponic normal (pH 6.5) and acidic (pH 4.0) conditions. (a) leaf SOD (U/g FW), (b) leaf POD ((U/g FW), (c) leaf CAT (U/g FW), (d) leaf APX (U/g FW), (e) root SOD (U/g FW), (f) root POD ((U/g FW), (f) root CAT (U/g FW), (g) root APX (U/g FW). Statistical differences among treatments were analyzed using appropriate multiple comparison tests. ns indicates not significant, while **, ***, and **** denote significant differences at P < 0.01, P < 0.001, and P < 0.0001, respectively. Data in the figures are means of eight replications (n = 8) ± standard deviation (SD).

#### 3.4.1 Superoxide Dismutase (SOD) Activity

SOD activity showed a strong induction under acidic conditions (pH 4) compared with pH 6.5 across all treatments, indicating enhanced superoxide radical generation under acid stress. In control plants, SOD activity at pH 4 was substantially higher (60.79 U mg□¹ FW) than at pH 6.5 (18.70 U mg□¹ FW), reflecting a basal stress response. Application of melatonin significantly increased SOD activity in a dose-dependent manner, with 100 µM MT reaching 137.1 U mg□¹ FW, representing a ∼128% increase over control, suggesting that MT promotes superoxide detoxification capacity under severe acidic stress (Fig. 3a). In contrast, under pH 6.5, SOD activity was relatively low and showed only moderate induction by MT, with a peak at 0.5 µM MT followed by a decline at higher concentrations (Fig. 3a). This differential response indicates that melatonin enhances ROS-scavenging systems primarily under stress conditions, while under near-optimal pH it avoids excessive activation of antioxidant machinery, thereby maintaining redox balance.

In roots, SOD activity exhibited a similar but more pronounced response. At pH 4, melatonin increased SOD activity from 54.34 U mg□¹ FW (control) up to 119.39 U mg□¹ FW at 100 µM MT (∼126% increase). This strong induction indicates that MT substantially enhances superoxide detoxification in roots exposed to acidic stress (Fig. 3b). At pH 6.5, the increase was smaller but still significant (18.70 U mg□¹ FW control to 42.29 U mg□¹ FW at 0.5 µM MT, ∼126% increase), however, MT application still increased SOD relative to control, though to a lesser extent than under acidic conditions (Fig. 3b). These results suggest that MT-mediated regulation of SOD is stress-responsive and tissue-specific, with a stronger effect in roots facing direct exposure to low-pH environments.

#### 3.4.2 Peroxidase (POD) Activity

POD activity exhibited an opposite trend compared with SOD, particularly under acidic conditions. At pH 4, control plants showed the highest POD activity (175.53 U mg□¹ FW), reflecting stress-induced activation of cell wall-associated and vacuolar peroxidases, which are often involved in stress signaling, lignification, and ROS-mediated defense responses. Melatonin application led to a progressive decrease in POD activity, with 100 µM MT reducing activity to 29.24 U mg□¹ FW (∼83% reduction compared to control). This decline suggests that melatonin alleviates oxidative stress, thereby reducing the requirement for POD-mediated hydrogen peroxide consumption (Fig. 3c). Importantly, this reduction highlights the dual role of melatonin: while it enhances ROS scavenging through SOD, CAT, and APX, it simultaneously suppresses excessive POD activity that is often associated with oxidative damage, rigidification of cell walls, and growth inhibition. Under pH 6.5, POD activity was generally lower than at pH 4 and displayed a distinct pattern, with moderate increases at higher MT concentrations (Fig. 3c). This suggests that under non-stress conditions, POD may participate more in fine-tuning redox signaling rather than bulk ROS detoxification.

In roots, POD activity displayed a similar pattern. At pH 4, POD activity also decreased from 168.59 U mg□¹ FW in control to 35.63 U mg□¹ FW at 100 µM MT (∼79% reduction), mirroring the trend observed in leaves and supporting the role of melatonin in alleviating oxidative stress by reducing H_2_O_2_ accumulation (Fig. 3d). However, at pH 6.5, POD activity increased progressively with melatonin concentration, reaching near-control or higher levels at 100 µM MT (Fig. 3d). This contrasting behavior highlights the dual role of melatonin: under stress, it suppresses excessive ROS-scavenging enzyme activity by lowering ROS production, whereas under non-stress conditions, it maintains or enhances POD activity to support redox homeostasis and signaling.

#### 3.4.3 Catalase (CAT) Activity

CAT activity was strongly stimulated by melatonin under both pH conditions, though the magnitude and pattern of induction differed. Under acidic pH, CAT activity increased from 139.4 U mg□¹ FW in control to 273.2 U mg□¹ FW at100 µM MT (∼96% increase over control) (Fig. 3e). This indicates that melatonin enhances the rapid decomposition of hydrogen peroxide into water and oxygen, a crucial mechanism for preventing oxidative damage under acid-induced stress. At pH 6.5, CAT activity was generally higher than control and showed a pronounced induction at low to moderate MT concentrations, with the highest activity observed at 0.5 µM MT, followed by a gradual decline at higher doses (Fig. 3e). This suggests that under near-neutral pH, lower MT concentrations are sufficient to optimize hydrogen peroxide homeostasis, while excessive MT may reduce the need for CAT due to improved redox equilibrium.

In roots, CAT activity similarly increased under acidic stress, from 167.6 U mg□¹ FW (control) to 331.6 U mg□¹ FW at 100 µM MT (∼98% increase) (Fig. 3f). At near-neutral pH, control activity was 147.9 U mg□¹ FW, and melatonin slightly increased CAT at low concentrations (0.5-5 µM MT) but decreased at higher doses (100 µM MT, 147.2 U mg□¹ FW) (Fig. 3f). These results indicate that melatonin effectively boosts CAT-mediated H_2_O_2_ scavenging, especially under acid stress, enhancing antioxidant defense in both leaves and roots.

#### 3.4.4 Ascorbate Peroxidase (APX) Activity

APX activity increased consistently with melatonin concentration under pH 4, indicating a strong MT-mediated activation of the ascorbate-glutathione cycle under acidic stress. At pH 4, control leaves had 28.8 U mg□¹ FW, which rose to 66.9 U mg□¹ FW at 100 µM MT (∼132% increase) (Fig. 3g). The highest APX activity at 100 µM MT suggests that APX plays a central role in fine-scale hydrogen peroxide detoxification in chloroplasts and cytosol when plants experience acid-induced oxidative pressure. Under pH 6.5, APX activity showed a different pattern, increased from 26.8 U mg□¹ FW (control) to 62.9 U mg□¹ FW at 0.5 µM MT, while higher MT concentrations showed slightly reduced activity (∼34.7–51.8 U mg□¹ FW), indicating a dose-dependent modulation (Fig. 3g). This response implies that under optimal conditions, melatonin primarily enhances APX activity at low doses to maintain signaling-compatible ROS levels, while higher concentrations may downregulate APX as oxidative stress is already minimal.

APX activity in roots increased significantly with MT under acidic conditions, from 18.3 U mg□¹ FW in control to 63.3 U mg□¹ FW at 100 µM MT (Fig. 3h). At pH 6.5, APX activity peaked at moderate MT doses (66.2 U mg□¹ FW at 0.5 µM), while higher MT doses slightly reduced activity (Fig. 3h), showing that APX is finely regulated by both MT concentration and pH to balance ROS detoxification with normal metabolism.

Collectively, these results demonstrate that melatonin orchestrates a coordinated antioxidant response in peanut leaves by enhancing ROS-scavenging enzymes (SOD, CAT, APX) while downregulating stress-associated POD activity, particularly under acidic conditions. This dual regulatory role allows melatonin to mitigate oxidative damage while preserving ROS signaling necessary for growth and adaptation. The contrasting responses observed between pH 4 and pH 6.5 further highlight that melatonin acts as a stress-dependent modulator of redox homeostasis rather than a constitutive antioxidant, enabling peanut plants to adapt efficiently to acidic stress environments.

### 3.5 Melatonin reduces hydrogen peroxide and lipid peroxidation contents in peanut under controlled hydroponic acidic conditions

Melatonin is widely recognized for its role in mitigating oxidative stress across various biological systems, including plants. We conducted a comprehensive study to assess how melatonin influences peanut seedling growth under acidic conditions. Our findings revealed that acid-treated plants exhibited significantly elevated levels of H_2_O_2_ and MDA compared to MT treated plants (Figs. 4a-d)

**Figure 4.**
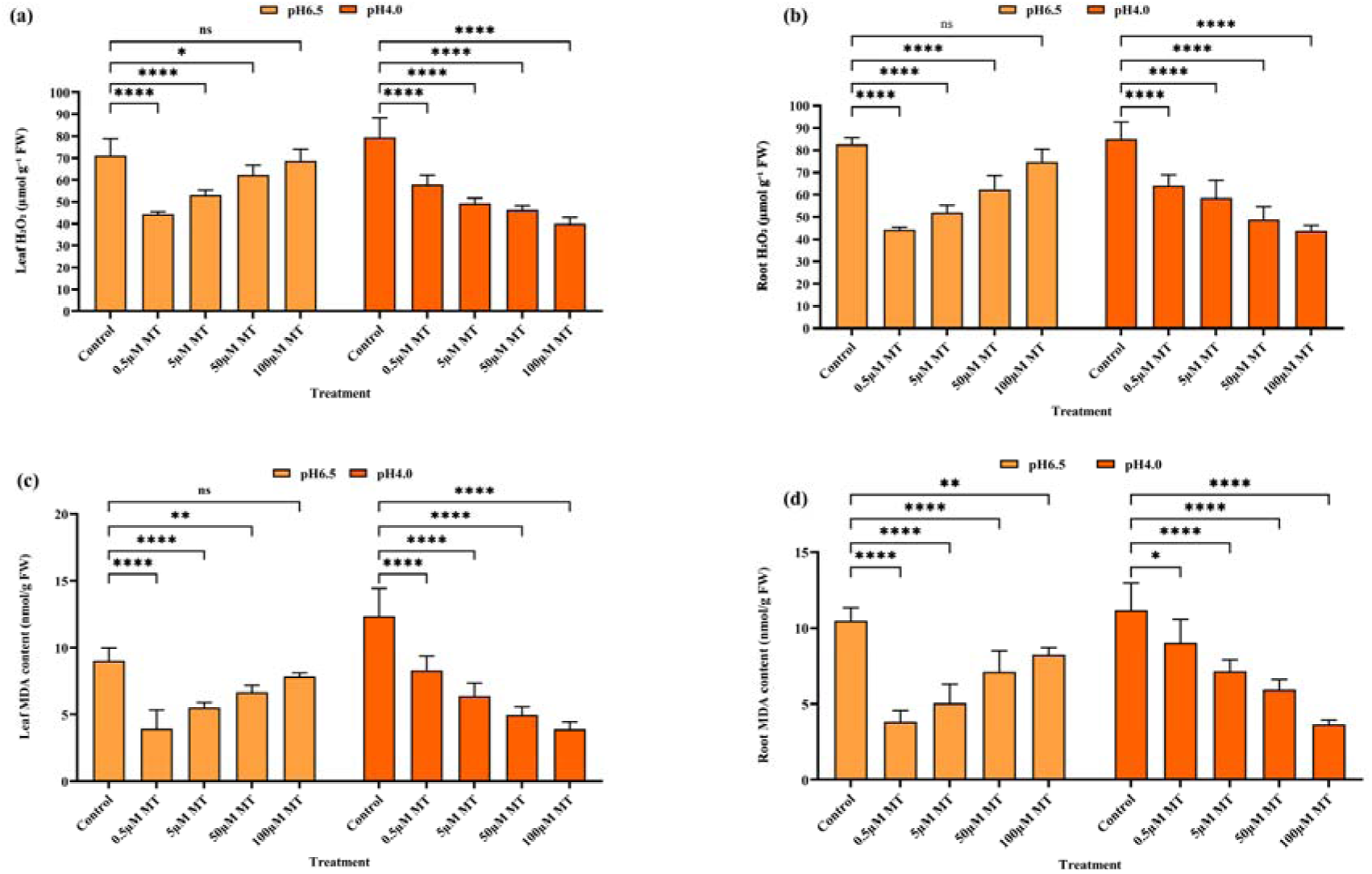
Effect of melatonin (MT) on H_2_O_2_ and MDA contents of peanut under control hydroponic normal (pH 6.5) and acidic (pH 4.0) conditions. (a) leaf H_2_O_2_ (µmol/g FW), (b) leaf MDA (nmol/g FW), (c) root H_2_O_2_ (µmol/g FW), (d) root MDA (nmol/g FW). Statistical differences among treatments were analyzed using appropriate multiple comparison tests. ns indicates not significant, while *, **, and **** denote significant differences at P < 0.05, P < 0.01, and P < 0.0001, respectively. Data in the figures are means of eight replications (n = 8) ± standard deviation (SD).

#### 3.5.1 Hydrogen Peroxide (H_2_O_2_) Content

Under control conditions (CK), H_2_O_2_ levels were high at pH 4 (79.40 µmol g□¹ FW) and pH 6.5 (71.06 µmol g□¹ FW), indicating substantial oxidative stress, particularly under acidic conditions. Melatonin application significantly reduced H_2_O_2_ accumulation at pH 4 in a dose-dependent manner. Compared with the control, 0.5 µM MT reduced H_2_O_2_ to 57.95 µmol g□¹ FW (≈28% decrease), 5 µM MT to 49.28 µmol g□¹ FW (≈38% decrease), 50 µM MT to 46.31 µmol g□¹ FW (≈43% decrease), and 100 µM MT to 39.99 µmol g□¹ FW (≈50% decrease) (Fig. 4a). In contrast, at pH 6.5, low MT concentrations initially decreased H_2_O_2_, with 0.5 µM MT reducing levels to 44.38 µmol g□¹ FW (≈36% decrease) and 5 µM MT to 53.03 µmol g□¹ FW (≈26% decrease). However, higher MT doses led to a reversal of this trend, with H_2_O_2_ increasing to 62.34 µmol g□¹ FW at 50 µM MT (≈11% decrease) and 68.71 µmol g□¹ FW at 100 µM MT (nearly comparable to control) (Fig. 4a). This suggests a dose and pH-dependent dual role of melatonin, acting as a strong antioxidant under acidic stress but allowing moderate ROS signaling at optimal pH.

Root tissues showed a similar but more pronounced response. Under pH 4, control plants accumulated 85.04 µmol g□¹ FW H_2_O_2_. Melatonin significantly lowered H□O□ to 64.25 µmol g□¹ FW at 0.5 µM (≈24% reduction), 58.54 µmol g□¹ FW at 5 µM (≈32% reduction), 48.76 µmol g□¹ FW at 50 µM (≈44% reduction), and 43.81 µmol g□¹ FW at 100 µM MT (≈51% reduction) (Fig. 4b). At pH 6.5, H□O□ content in roots declined at low MT doses (44.25 µmol g□¹ FW at 0.5 µM; ≈46% decrease) but increased at higher concentrations, reaching 74.78 µmol g□¹ FW at 100 µM MT (≈9% decrease compared with control) (Fig. 4b). These results indicate that melatonin efficiently scavenges excessive ROS under acid stress, while at neutral pH high MT may permit ROS accumulation for signaling functions.

#### 3.5.2 Malondialdehyde (MDA) Content

MDA content, an indicator of lipid peroxidation, was markedly higher under acidic conditions. In leaves at pH 4, control plants exhibited 12.35 nmol g□¹ FW MDA. Melatonin treatments progressively reduced MDA levels to 8.29 nmol g□¹ FW at 0.5 µM (≈32% reduction), 6.38 nmol g□¹ FW at 5 µM (≈48% reduction), 4.94 nmol g□¹ FW at 50 µM (≈60% reduction), and 3.89 nmol g□¹ FW at 100 µM MT (≈69% reduction) (Fig. 4c). At pH 6.5, control MDA content was lower (9.0 nmol g□¹ FW). □ow MT concentrations further decreased MDA to 3.92 nmol g□¹ FW at 0.5 µM (≈56% reduction) and 5.52 nmol g□¹ FW at 5 µM (≈39% reduction). However, higher MT doses increased MDA to 6.66 nmol g□¹ FW at 50 µM (≈26% reduction) and 7.83 nmol g□¹ FW at 100 µM MT (≈12% reduction) (Fig. 4c), again reflecting melatonin’s concentration-dependent dual behavior.

Root MDA followed trends similar to leaves but with overall lower values. At pH 4, control roots had 11.16 nmol g□¹ FW MDA. This decreased to 9.04 nmol g□¹ FW at 0.5 µM (≈19% reduction), 7.15 nmol g□¹ FW at 5 µM (≈35% reduction), 5.93 nmol g□¹ FW at 50 µM (≈46% reduction), and 3.65 nmol g□¹ FW at 100 µM MT (≈68% reduction) (Fig. 4d). At pH 6.5, MDA declined sharply at low MT doses (3.83 nmol g□¹ FW at 0.5 µM; ≈64% reduction) but increased again at higher concentrations, reaching 8.25 nmol g□¹ FW at 100 µM MT (≈20% reduction compared with control) (Fig. 4d). This indicates that melatonin effectively protects membrane integrity under acidic stress, while excessive doses at optimal pH may attenuate this protective effect.

Collectively, these results demonstrate that melatonin markedly alleviates oxidative stress and lipid peroxidation in peanut leaves and roots, particularly under acidic (pH 4) conditions. The strong reduction in H_2_O_2_ and MDA at pH 4 indicates that melatonin primarily functions as an antioxidant under stress. In contrast, under pH 6.5, low melatonin concentrations enhance oxidative protection, whereas higher doses partially restore ROS and lipid peroxidation levels, suggesting a regulated balance between ROS scavenging and signaling. This dual role of melatonin highlights its importance in fine-tuning redox homeostasis in peanut plants under contrasting pH environments.

### 3.6 Melatonin induces dose-dependent upregulation of HD-ATPase genes under controlled hydroponic acid stress

To investigate the transcriptional response of plasma membrane H□-ATPase genes to melatonin (MT) under acidic conditions, the relative expression levels of *AhAH1* and *AhAH2* were analyzed in peanut leaves and roots under acid stress (Figs. 5a and b). Both genes are central to proton extrusion and pH homeostasis during environmental stress.

**Figure 5.**
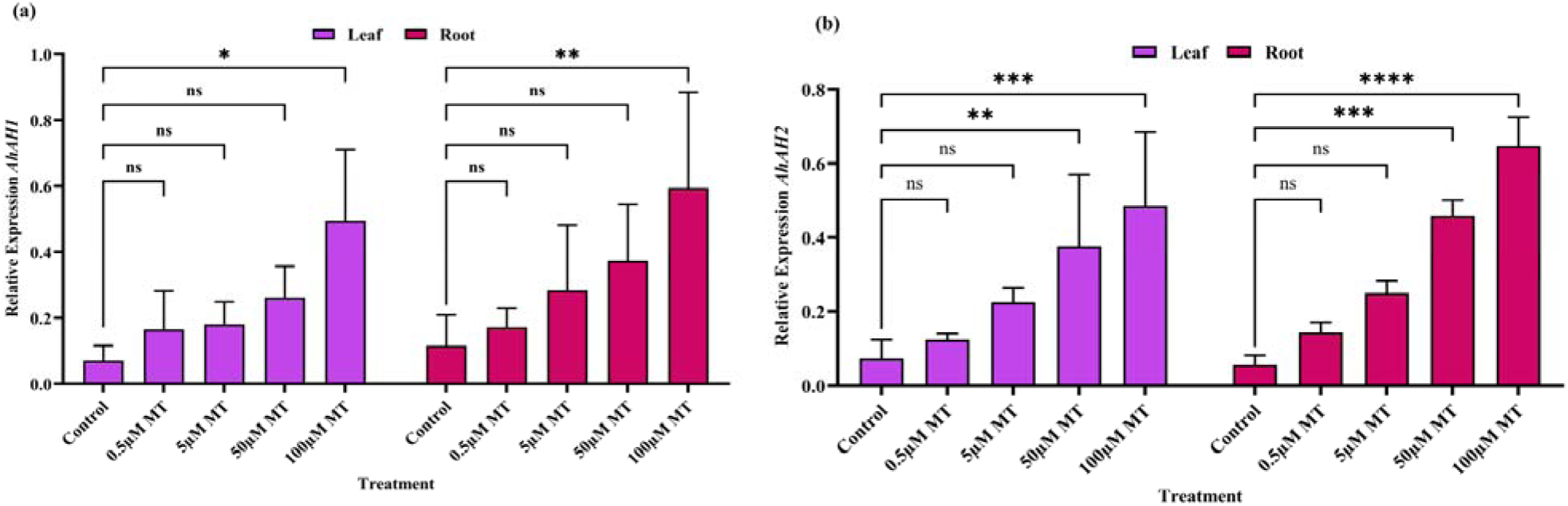
Melatonin-induced expression of plasma membrane HD-ATPase genes in peanut leaves and roots under acid conditions. (a) Relative expression of *AhAH1* and (b) relative expression of *AhAH2* in leaves and roots under different melatonin (MT) concentrations (0.5, 5, 50, and 100 µM) compared with the control. Bars represent mean ± SD of biological replicates. Statistical differences among treatments were analyzed using appropriate multiple comparison tests. ns indicates not significant, while *, **, ***, and **** denote significant differences at P < 0.05, P < 0.01, P < 0.001, and P < 0.0001, respectively.

Melatonin treatment significantly enhanced the expression of plasma membrane H□-ATPase genes *AhAH1* and *AhAH2* in peanut seedlings, with a pronounced induction under acidic conditions. The basal expression of *AhAH1* was low in control plants, with minimal transcript accumulation observed in both leaves and roots. MT supplementation resulted in a consistent and progressive increase in *AhAH1* expression at all concentrations tested. At 0.5 and 5 µM MT, *AhAH1* transcript levels increased by approximately 2.4, and 2.6-fold in leaves and 1.5, and 2.5-fold in roots relative to the control (Fig. 5a). This induction intensified with increasing MT concentration, reaching ∼3.7-fold in leaves and ∼3.2-fold in roots at 50 µM MT. The strongest response was observed at 100 µM MT, where *AhAH1* expression increased by ∼7.0-fold in leaves and ∼5.1-fold in roots compared with control plants, indicating robust, dose-dependent transcriptional activation (Fig. 5a). The stronger induction in roots highlights the pivotal role of root-localized H□-ATPase activity in coping with acidic stress.

A similar positive response was observed for *AhAH2*, which exhibited higher basal expression than *AhAH1* in control plants, particularly in roots. Nevertheless, MT treatment further enhanced *AhAH2* expression at all concentrations under acid stress. At low MT concentrations (0.5 and 5 µM), *AhAH2* transcript abundance increased by approximately 1.7- and 3.1-fold in leaves and 2.6- and 4.5-fold in roots, respectively, relative to the control (Fig. 5b). This upregulation became progressively stronger with increasing MT dosage, reaching approximately 5.1-fold in leaves and 8.2-fold in roots at 50 µM MT. The maximum induction occurred at 100 µM MT, where *AhAH2* expression was elevated by approximately 6.6-fold in leaves and 11.6-fold in roots compared with control plants (Fig. 5b), demonstrating a robust and concentration-dependent transcriptional response. Importantly, All MT treatments significantly increased *AhAH2* transcript levels relative to the control, with the highest induction observed at 100 µM MT. This consistent upregulation under acidic stress suggests that *AhAH2* is positively responsive to melatonin and likely contributes to enhanced proton extrusion and pH homeostasis. The pronounced upregulation of *AhAH2* at higher melatonin concentrations suggests that this gene encodes a stress-adaptive H□-ATPase that remains transcriptionally active under strong melatonin stimulation, thereby contributing to the maintenance of cytosolic pH homeostasis and sustained root function under acidic stress conditions.

Overall, these results demonstrate that melatonin induces a consistent, concentration-dependent increase in H□-ATPase gene expression at all MT treatments relative to the control, with the strongest fold induction occurring under acidic conditions. This transcriptional enhancement provides molecular evidence for melatonin-mediated reinforcement of proton transport capacity, supporting improved acid tolerance in peanut seedlings.

### 3.7 Field Soil Validation of Melatonin Effects under Natural Acidic Conditions

The pot experiment conducted using naturally acidic soil (pH 4.3-4.5) collected from the southern region of Henan Province demonstrated growth inhibition patterns comparable to those observed under hydroponic acidic conditions. Control plants grown without melatonin exhibited reduced germination percentage, shorter root and shoot lengths, and significantly lower fresh and dry biomass, indicating the detrimental impact of natural soil acidity on peanut growth (Fig. 6). Melatonin application showed a clear dose-dependent response, with 5 µM MT producing the most pronounced improvement in early seedling establishment and vegetative growth parameters. At this concentration, root elongation, shoot height, and biomass accumulation increased markedly compared with the untreated control, whereas higher concentrations (50 and 100 µM) resulted in a gradual decline in growth performance, suggesting an optimal threshold for melatonin efficacy under soil conditions (Fig. 6).

**Figure 6.**
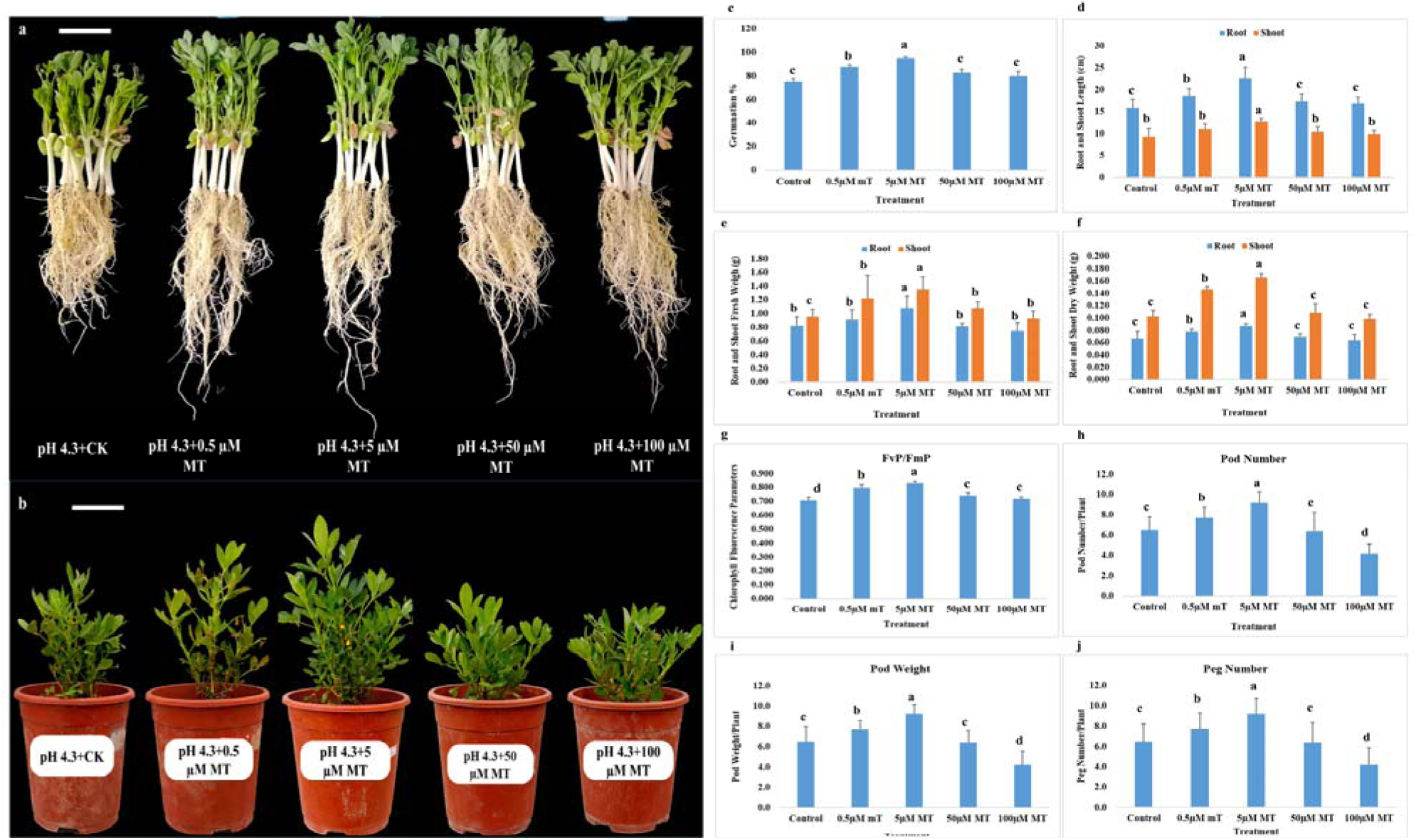
Effect of melatonin (MT) on seed germination and seedling growth, chlorophyll fluorescence and yield-related traits under natural acidic field-soil (pH 4.3-4.5) conditions on peanut. **(a)** seedling growth phenotypes, **(b)** maturity stage phenotypes, **(c)** seed germination (%), **(d)** root and shoot length (cm), **(e)** root and shoot fresh weight (g), **(f)** root and shoot dry weight (g), **(g)** chlorophyll fluorescence (FvP/FmP) **(h**) pod number, **(i)** pod weight (g), **(j)** peg number. Statistical differences among treatments were analyzed using appropriate multiple comparison tests. Different lowercase letters above bars show significant differences (p < 0.05) between CK and MT concentrations. Data in the figures are means of 10 replications (n = 10) ± standard deviation (SD). Bar = 2 cm.

Physiological assessment revealed that chlorophyll fluorescence (Fv/Fm) values were significantly higher in plants treated with 5 µM MT, reflecting enhanced photosystem II efficiency and reduced photoinhibition under acidic stress (Fig. 6). Yield-related traits followed a similar trend, where 5 µM MT treatment led to increased peg number per plant, greater pod number, and higher total pod weight relative to both the control and higher MT concentrations (Fig. 6). Collectively, these findings indicate that melatonin confers protective effects against natural soil acidity in a concentration-dependent manner, with moderate doses maximizing plant growth, physiological stability, and reproductive performance. The consistency between hydroponic and soil-based observations confirms that melatonin-mediated acid stress tolerance is reproducible in realistic agricultural environments, while also highlighting the importance of dose optimization under complex soil conditions.

## 4. Discussion

### 4.1 Melatonin alleviates acid stress-induced growth inhibition by promoting seed germination and seedling growth in peanut

Acidic stress conditions severely constrain seed germination and early seedling establishment by disrupting metabolic activation, reserve mobilization, and cell expansion (Msimbira & Smith, 2020; Hasan et al., 2025). Seed germination and early seedling establishment are among the most pH-sensitive developmental stages in crops, and acidic stress is known to impair metabolic activation, reserve mobilization, and cell expansion (Shavrukov & Hirai, 2016; Ali *et al*., 2023; Wang *et al*., 2025b). In this study, peanut seeds exposed to acidic pH (4.0) showed a clear reduction in germination percentage and seedling growth compared with near-neutral pH (6.5), confirming the high pH sensitivity of early developmental stages. Exogenous melatonin (MT) markedly alleviated this inhibition, significantly improving germination and promoting vigorous seedling growth, with a much stronger relative effect under acidic stress than under near-neutral conditions (Figs.1a-h). The enhanced germination under acidic pH likely reflects improved metabolic activation during imbibition, as MT is known to stimulate hydrolytic enzymes such as α-amylases and proteases, thereby facilitating efficient mobilization of stored reserves under stress (iu *et al*., 2023; Fu *et al*., 2025). In addition, MT interacts with key phytohormones, including gibberellins and auxins, which regulate radicle protrusion and early seedling vigor, further supporting rapid germination under low-pH conditions (Arnao & Hernández-Ruiz, 2019; Fu et al., 2025). Consistent with improved germination, MT substantially enhanced root and shoot growth and biomass accumulation under acidic conditions, indicating improved seedling establishment (Figs. 1a-h). Similar MT-mediated promotion of early growth under unfavourable soil conditions has been reported in other crops, where MT improved root development and stress resilience by maintaining cellular homeostasis and nutrient acquisition (Muhammad *et al*., 2024; Chung *et al*., 2025; Hasan et al., 2025). In contrast, under near-neutral pH, low MT doses were sufficient to slightly enhance seedling performance, whereas higher concentrations often reduced growth and physiological indices, highlighting the role of MT as a stress-responsive regulator rather than a constitutive growth promoter. This inhibitory effect likely results from the disruption of finely tuned redox and hormonal signaling, as melatonin also functions as a regulatory molecule and excessive levels can suppress seedling growth under non-stress conditions (Arnao & Hernández-Ruiz, 2019; Ameen *et al*., 2024; Hasan et al., 2025). Collectively, these results demonstrate that melatonin effectively mitigates acid-induced inhibition of peanut seed germination and early seedling growth, underscoring its potential for improving crop establishment in acidic soils.

### 4.2 Melatonin enhances photosynthetic pigment stability and efficiency under acid stress in peanut

Acidic stress markedly impaired photosynthetic pigment accumulation in peanut seedlings, as evidenced by reduced chlorophyll a, chlorophyll b, total chlorophyll, and carotenoid contents under pH 4, indicating disrupted chloroplast function and pigment biosynthesis (Figs. 2a-d). Such reductions are well documented consequences of low pH-induced proton toxicity, metal ion imbalance, and oxidative damage, which collectively destabilize chlorophyll synthesis and accelerate pigment degradation during early vegetative growth (Sharma *et al*., 2020; Yang *et al*., 2023; Jiang *et al*., 2025b). Our results showed that exogenous melatonin application substantially alleviated this inhibition, resulting in a dose-dependent restoration of chlorophyll and carotenoid levels under acidic conditions, with maximal pigment accumulation observed at higher MT concentrations (Figs. 2a-d). This response suggests that melatonin preserves pigment integrity under acid stress, likely by protecting chloroplast membranes and photosynthetic complexes from oxidative injury through its strong antioxidant and redox-buffering capacity (Dou et al., 2025; Hasan et al., 2025). Carotenoid accumulation followed a similar pattern, increasing progressively with MT concentration under pH 4, suggesting enhanced photoprotection and reactive oxygen species scavenging (Figs. 2a-d). The associated increase in carotenoid content under acidic conditions further supports a protective role of melatonin, as carotenoids are essential for quenching excess excitation energy and preventing photo-oxidative damage under stress. Similar MT-induced stabilization of photosynthetic pigments has been reported in crops exposed to salinity, drought, and metal toxicity, where melatonin maintained chlorophyll biosynthesis and delayed stress-induced senescence (An *et al*., 2025; Basharat et al., 2025; Chen *et al*., 2025a; Hasan et al., 2025; Li *et al*., 2025e). In contrast, under near-neutral pH (6.5), pigment levels were inherently high in control plants, and low MT concentrations were sufficient to enhance chlorophyll accumulation, whereas higher doses led to pigment stabilization or decline. This concentration-dependent response indicates that melatonin primarily functions as a stress-responsive regulator, exerting its strongest effects when pigment homeostasis is challenged rather than under optimal growth conditions (Ali et al., 2023; Yang et al., 2023). Overall, these results demonstrate that melatonin enhances photosynthetic pigment stability in peanut in a pH and dose-dependent manner, with higher MT concentrations effectively counteracting acid-induced pigment loss. This protective role of melatonin on chlorophyll and carotenoid metabolism likely contributes to improved photosynthetic competence and stress resilience of peanut seedlings under acidic environments, a condition that remains underexplored in crop systems.

### 4.3 Melatonin-regulated antioxidant redox homeostasis confers acid stress tolerance in peanut

Acidic stress markedly disrupts redox homeostasis in plants by enhancing reactive oxygen species (ROS) production and impairing antioxidant efficiency, thereby constraining early growth and physiological stability (Hlihor *et al*., 2022; Jahan *et al*., 2022; Hasan et al., 2025). In the present study, peanut seedlings exposed to acidic pH (4) exhibited strong oxidative pressure, as evidenced by differentially modulated basal activities of stress-responsive antioxidant enzymes, such as SOD, POD, CAT and APX. Exogenous MT application significantly reprogrammed the antioxidant defense system, resulting in a coordinated and stress-adaptive response. Under acidic conditions, MT markedly enhanced SOD, CAT, and APX activities in both leaves and roots, indicating improved superoxide dismutation and hydrogen peroxide detoxification, while simultaneously suppressing excessive POD activity (Figs. 3a-h). The marked reduction in POD activity observed in melatonin-treated peanut leaves and roots under acidic conditions likely reflects melatonin-mediated alleviation of oxidative stress rather than suppression of antioxidant capacity. Exogenous melatonin significantly reduced POD activity in a dose-dependent manner, indicating that MT effectively lowers ROS production and H_2_O_2_ availability, thereby reducing the requirement for POD-dependent scavenging (Figs. 3b and c). It has been reported that melatonin has been shown to modulate antioxidant enzyme activities and reduce ROS accumulation under various stress conditions, contributing to redox homeostasis (meta-analysis in rice and other plants) (Yang *et al*., 2021; Muhammad *et al*., 2025). Moreover, in other systems melatonin reduced ROS accumulation while lowering peroxidase activity, supporting the interpretation that decreased POD activity can accompany improved oxidative balance (Yang et al., 2021; Hasan et al., 2025). Thus, the reduced POD activity in MT-treated peanut seedlings reflects a functional shift from stress-induced peroxidase activation to a more energy-efficient antioxidant strategy, highlighting melatonin’s dual role as both a ROS scavenger and a redox regulator under acidic stress (Figs. 3b and c). This pattern suggests that MT not only strengthens ROS-scavenging capacity but also limits stress-associated peroxidative reactions linked to cell wall rigidification and growth inhibition. Similar MT-mediated enhancement of enzymatic antioxidants and redox balance under acidic or oxidative stress has been reported in many crops, where MT improved stress tolerance by activating ROS-detoxifying enzymes and their associated genes (Ahammed *et al*., 2020; Ahammed *et al*., 2024; Chakraborty & Raychaudhuri, 2025; Chung et al., 2025; Li *et al*., 2025c). Notably, the contrasting responses observed at pH 6.5 further support the stress-dependent role of melatonin. Under near-neutral conditions, MT induced only moderate or dose-limited changes in antioxidant activity, indicating that MT fine-tunes redox signaling rather than overstimulating defense pathways when oxidative stress is minimal (Figs. 3a-h). This dual behavior, strong activation under stress and restrained modulation under optimal conditions, has been recognized as a hallmark of melatonin function in plants (Moustafa-Farag *et al*., 2020; Menhas *et al*., 2025; Shehzadi *et al*., 2025). Collectively, these findings demonstrate that melatonin enhances acid-stress tolerance in peanut by orchestrating a balanced antioxidant response, promoting efficient ROS detoxification while preserving redox signaling, thereby enabling physiological stability and growth under low-pH environments.

### 4.4 Melatonin fine-tunes redox homeostasis by limiting hydrogen peroxide accumulation and membrane lipid peroxidation under acidic stress

The pronounced accumulation of hydrogen peroxide (H_2_O_2_) and malondialdehyde (MDA) in peanut seedlings under acidic conditions confirms that low pH imposes severe oxidative stress, leading to excessive ROS generation and membrane lipid peroxidation (Hlihor et al., 2022; Hasan et al., 2025). Under pH 4, control plants exhibited markedly elevated H_2_O_2_ and MDA levels in both leaves and roots, reflecting impaired redox balance and enhanced oxidative damage. Exogenous melatonin significantly and dose-dependently reduced H_2_O_2_ accumulation under acidic conditions, with the highest MT concentration lowering H_2_O_2_ by ∼50% in leaves and ∼51% in roots relative to the control (Figs. 4a and b). This strong suppression of ROS indicates that melatonin acts primarily as an efficient antioxidant under acid stress, either through direct free-radical scavenging or by reinforcing enzymatic detoxification pathways, thereby reducing cellular H_2_O_2_ availability (Arnao & Hernández-Ruiz, 2019; Ahammed et al., 2020; Ahammed et al., 2024; Hasan et al., 2025; Muhammad et al., 2025).

Consistent with reduced ROS levels, MDA content declined sharply in MT-treated seedlings under pH 4, with up to ∼69% and ∼68% reductions in leaves and roots, respectively, indicating effective protection of membrane integrity (Figs. 4c and d). The tight coupling between decreased H_2_O_2_ and lower lipid peroxidation suggests that melatonin limits oxidative damage by stabilizing cellular membranes and preventing ROS-induced fatty acid oxidation, a response widely associated with improved stress tolerance (Ali et al., 2023; Ahammed et al., 2024; Hasan et al., 2025). In contrast, under near-neutral pH (6.5), low MT concentrations enhanced oxidative protection, whereas higher doses partially restored H_2_O_2_ and MDA levels (Figs. 4a-d), supporting the concept that melatonin permits controlled ROS accumulation for signaling under non-stress conditions (Hasan et al., 2025; Li *et al*., 2025a). Collectively, these results demonstrate that melatonin functions as a redox modulator in peanut, acting predominantly as an antioxidant under acidic stress while maintaining ROS-dependent signaling at optimal pH, thereby ensuring balanced cellular homeostasis.

### 4.5 Melatonin enhances plasma membrane H**D**-ATPase transcription to maintain pH homeostasis under acidic stress

Plasma membrane H□-ATPases are key regulators of proton extrusion, cytosolic pH stability, and nutrient uptake, and their function becomes especially critical under acidic stress, where excessive proton influx disrupts cellular metabolism and root growth (Chao & Chao, 2025; Dong *et al*., 2025). In the present study, melatonin (MT) significantly and dose-dependently upregulated the expression of the H□-ATPase genes *AhAH1* and *AhAH2* in peanut seedlings under acidic conditions, with markedly stronger induction in roots than in leaves. This transcriptional response provides molecular evidence that MT reinforces proton transport to the apoplast to improve acid tolerance (Figs. 5a and b).

The strong MT-induced expression of *AhAH1* and *AhAH2* is consistent with previous reports showing that melatonin enhances H□-ATPase activity and expression under various abiotic stresses (Wang *et al*., 2025c). For example, MT has been shown to stimulate plasma membrane H□-ATPases in cucumber and tomato under salinity and nutrient stress, thereby stabilizing intracellular pH and improving ion homeostasis (Li et al., 2022; Zeng et al., 2024; Li *et al*., 2025b). These findings align closely with our observation that *AhAH1* and *AhAH2* expression was most strongly induced at higher melatonin concentrations under pH 4, highlighting a conserved role of melatonin in activating proton transport machinery under acidic stress (Figs. 5a and b). Under acidic or aluminium-stress conditions, enhanced H□-ATPase activity promotes proton efflux and mitigates rhizotoxic effects, supporting root elongation and function (Zhang *et al*., 2017; Munyaneza *et al*., 2024). The robust upregulation of *AhAH2* at higher melatonin doses indicates that this isoform may function as a stress-adaptive proton pump, supporting sustained root activity, membrane potential stability, and nutrient acquisition under acidic conditions. The pronounced induction of *AhAH2*, particularly in roots, suggests isoform-specific stress responsiveness, in line with studies in Arabidopsis and rice, where certain H□-ATPase genes are preferentially activated under low-pH stress to maintain root apoplastic and cytosolic pH balance (Gámez-Arjona *et al*., 2022; Chakraborty *et al*., 2024; Jain & Schmidt, 2024).

Mechanistically, melatonin-mediated upregulation of H□-ATPase genes likely reflects its integrative role in redox and hormonal signaling. Melatonin modulates ROS levels and interacts with auxin and calcium-dependent pathways, both of which are known regulators of H□-ATPase transcription and activation (Huang et al., 2022; Yang *et al*., 2024; Chung et al., 2025). By alleviating oxidative stress while maintaining controlled ROS signaling, MT may create favorable cellular conditions for sustained proton pump expression and activity under acidic environments. Overall, these findings demonstrate that melatonin enhances acid stress tolerance in peanut by transcriptionally activating plasma membrane H□-ATPase genes, particularly *AhAH2* in roots. This response likely strengthens proton extrusion, stabilizes intracellular pH, and supports root performance under low-pH conditions, highlighting H□-ATPase regulation as a key molecular component of melatonin-mediated adaptation to acidic stress in crops.

### 4.6 Field Soil Validation and Dose-Dependent Response of Melatonin

The soil-based validation experiment verifies the ecological applicability of melatonin-mediated acid stress tolerance previously observed under hydroponic conditions. As illustrated in Figure 6, peanut plants cultivated in naturally acidic soil (pH 4.3-4.5) exhibited substantial growth suppression in the untreated control, whereas melatonin supplementation significantly improved seedling establishment and vegetative development. Notably, the superior performance of the 5 µM MT treatment compared with higher concentrations confirms a concentration-dependent regulatory effect of melatonin under complex soil environments. This biphasic or hormetic response is consistent with earlier studies reporting that moderate melatonin levels stimulate plant growth while excessive concentrations may inhibit physiological processes (Arnao & Hernández-Ruiz, 2018; Sun *et al*., 2021; Wang *et al*., 2022). Comparable dose-specific enhancements in root elongation and biomass accumulation have been documented in maize and soybean under abiotic stress, suggesting that melatonin functions as a broadly conserved plant biostimulant (Ahmad *et al*., 2021; Hasan et al., 2025; He *et al*., 2025; Zhao *et al*., 2025).

Physiological stability under acidic stress was further supported by chlorophyll fluorescence analysis, in which elevated Fv/Fm ratios in the 5 µM treatment (see Figure Y) indicate preserved photosystem II efficiency and reduced photoinhibition. Similar protective effects of melatonin on photosynthetic machinery have been widely reported under drought, salinity, and heavy-metal stress conditions (Chen *et al*., 2025b; Moradi *et al*., 2026). Importantly, the improved physiological performance translated into reproductive benefits, as reflected by increased peg number, pod number, and total pod weight per plant **(Fig. 6)**, reinforcing the agronomic significance of melatonin application. Parallel yield-related improvements have also been described in rice and wheat under environmental stress, confirming the broader crop relevance of melatonin-induced resilience (Yan *et al*., 2024; Li *et al*., 2025d).

Interestingly, the divergence in optimal melatonin concentration between hydroponic and soil systems observed in this study suggests that environmental context strongly influences melatonin bioactivity. Soil-specific factors such as adsorption to organic matter, microbial degradation, and heterogeneous nutrient availability likely reduce effective melatonin concentration and shift the optimal dose toward lower levels. Similar observations have been noted in soil-plant interaction studies, emphasizing the importance of dosage optimization under field-like conditions (Arnao & Hernández-Ruiz, 2019; Khattak *et al*., 2023; Aijaz *et al*., 2025). Collectively, the present results not only validate previous findings but also extend current understanding by demonstrating that melatonin efficacy is both environment-dependent and dose-specific, thereby highlighting the necessity of system-tailored application strategies for sustainable crop production in acid-affected soils.

### 4.7 Interaction between melatonin concentration and pH regulates peanut physiological responses

Across all measured parameters, including germination, seedling growth, biomass accumulation, photosynthetic pigments, oxidative stress markers, antioxidant enzyme activities, and H□-ATPase gene expression, a clear interaction between melatonin concentration and pH level was observed. Acidic conditions (pH 4) consistently amplified the positive effects of melatonin, whereas near-neutral pH (6.5) markedly reduced or altered MT responsiveness. This interaction indicates that melatonin does not act as a simple growth promoter, but rather functions as a stress-dependent regulator whose efficacy is tightly linked to environmental pH (Fig. 7).

**Figure 7.**
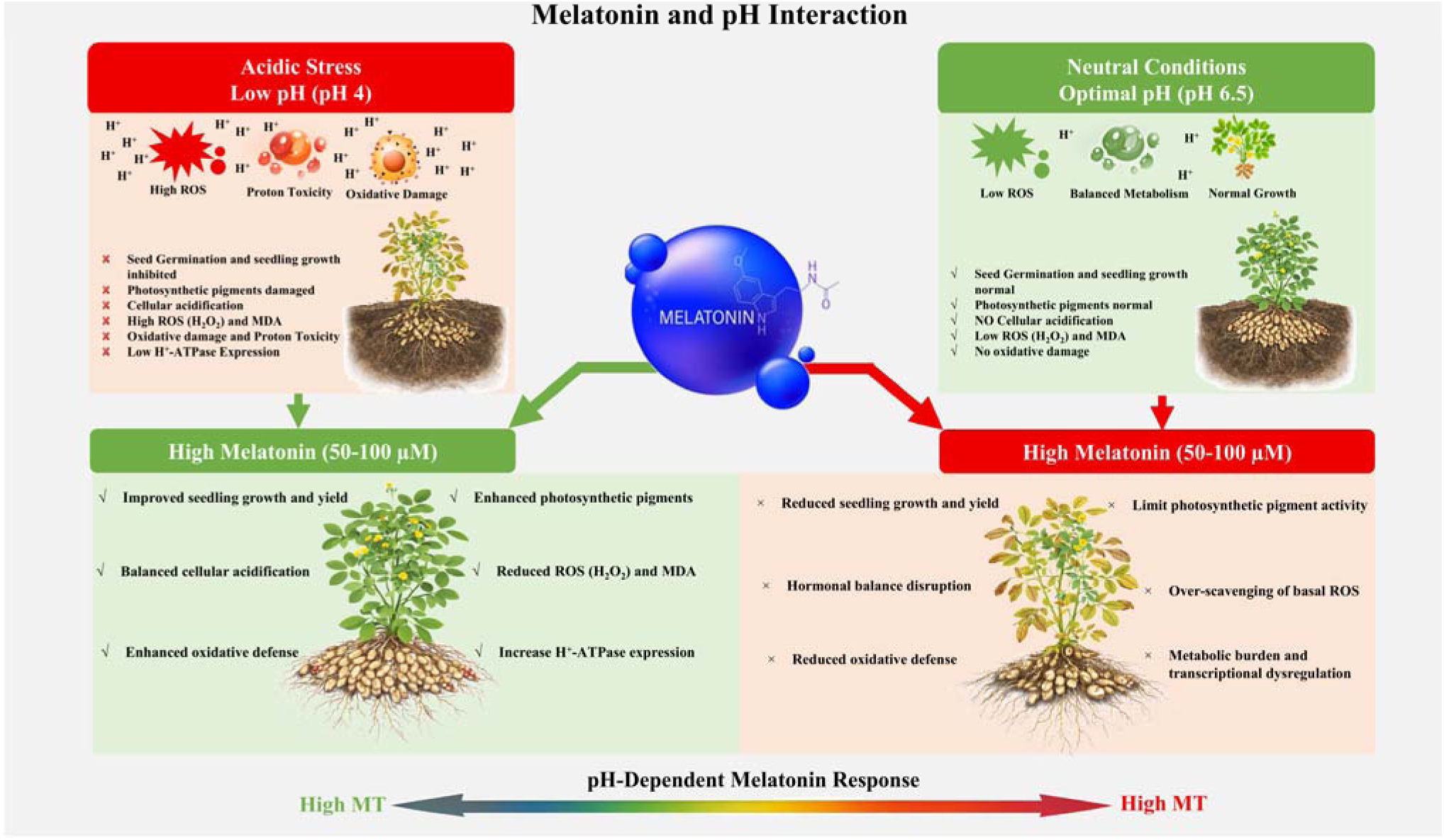
Melatonin and pH interaction shows that a higher melatonin level is required to alleviate seedling growth and yield, oxidative damage, photosynthetic pigments, ROS and HlJ-ATPase genes expression and pH imbalance under acidic stress, whereas excessive melatonin can disrupt normal growth under optimal conditions.

Under acidic stress, increasing MT concentrations progressively improved germination, root and shoot development, pigment accumulation, redox homeostasis, and proton pump gene expression. These responses reflect a coordinated mitigation of proton toxicity, oxidative damage, and metabolic disruption, suggesting that higher MT availability is required to counteract the elevated ROS burden and pH imbalance typical of acid stress (Fig. 7). Similar pH or stress-intensity-dependent melatonin responses have been reported under salinity, metal toxicity, and drought, where stronger stress necessitated higher MT doses to restore cellular homeostasis (Arnao & Hernández-Ruiz, 2019; Ahmad et al., 2023b; Ahammed et al., 2024; An et al., 2025; Basharat et al., 2025).

In contrast, under near-neutral pH, most physiological parameters peaked at low MT concentrations and declined at higher doses, indicating that excessive melatonin can disrupt optimal redox and hormonal signaling when stress pressure is minimal (Fig. 7). Basal ROS are essential signaling molecules for normal growth, photosynthesis, and development, and over-scavenging by high MT levels may suppress these signaling pathways, leading to reduced growth and metabolic efficiency (Back, 2021; Liu et al., 2023; Dzinyela *et al*., 2025). Thus, the MT and pH interaction demonstrates that melatonin acts within an optimal concentration window that shifts upward with increasing stress severity.

Collectively, this interaction highlights melatonin as a conditional bioregulator whose beneficial range depends on environmental pH, emphasizing the importance of dose optimization when applying MT in agronomic systems.

### Conclusion

The present study provides comprehensive evidence that exogenous melatonin application markedly enhances peanut tolerance to acidic stress by integrating redox regulation, growth maintenance, and pH homeostasis. Acidic conditions severely restricted peanut growth, pigment synthesis, and differentially affected antioxidants, while inducing excessive H_2_O_2_ accumulation and membrane lipid peroxidation. Melatonin application significantly reversed these effects, particularly at pH 4, by suppressing oxidative damage, protecting the photosynthetic apparatus, and sustaining biomass production. This protection was associated with a coordinated reprogramming of the antioxidant system, characterized by enhanced SOD, CAT and APX activities and reduced POD activity, indicating alleviation of oxidative pressure and a shift toward more efficient H_2_O_2_ detoxification. In addition, melatonin also strongly induced the plasma membrane H□-ATPase genes *AhAH1* and *AhAH2*, especially in roots, providing molecular evidence that melatonin reinforces proton extrusion capacity and cellular pH stability under acid stress.

Importantly, the soil-based validation experiment extended these mechanistic insights to realistic agricultural scenarios. Hydroponic experiments revealed that higher melatonin concentrations produced the strongest protective responses due to uniform nutrient availability and direct root exposure. In contrast, soil-based validation under naturally acidic field soil (pH 4.3-4.5) confirmed that moderate concentrations (5 µM) most effectively improved germination, vegetative growth, chlorophyll fluorescence (Fv/Fm), and yield-related traits, whereas excessive doses reduced performance. Collectively, these findings establish melatonin as a multifunctional and context-dependent bioregulator whose optimal concentration varies with environmental complexity. The integration of mechanistic hydroponic evidence with realistic soil validation provides both theoretical insight and practical guidance for optimizing melatonin application to enhance peanut establishment and productivity on acid-affected agricultural soils.

## Author Contributions

**Muhammad Hafeez Ullah Khan:** conceptualization, investigation, writing – original draft. **Fu Ruoyi:** methodology and formal analysis. **Ali Muhammad:** visualization and formal analysis. **Shuaichao Zheng:** data curation and formal analysis. **Daijing Zhang:** writing – review and editing. **Zhiyong Zhang:** supervision, funding acquisition, writing – review and editing. **Quanyong Liu:** supervision, funding acquisition, writing – review and editing.

## Funding and Acknowledgement

This research was partially supported by the National Natural Science Foundation of China (31571600, 31271648), the Key R&D and Promotion Projects of Henan Province (222102110303, 232102110221, 242102111094, 252102111072), the International Science and Technology Cooperation Project of Henan Province (242102521050), and the Henan Center of Outstanding Overseas Scientists (GZS2024018). We gratefully acknowledge Professor Xiaoping Pan from East Carolina University for her great input, comments, and proofreading of this paper.

## Data Availability Statement

The data that support the findings of this study are available from the corresponding author upon reasonable request.

